# HIV broadly neutralizing antibodies with enhanced Fc-CD16 affinity increase NK cell ADCC even with limited envelope binding

**DOI:** 10.64898/2026.04.27.721006

**Authors:** Claudia Melo, Teresa Murphy, Carissa S Holmberg, Meagan Kelly, Elyse K McMahon, Jonathan S Lochner, Rebecca M Lynch, Alberto Bosque

**Affiliations:** Department of, Microbiology, Immunology & Tropical Medicine, Washington, DC, USA., The George Washington University, Washington, DC, USA

**Keywords:** 1. NK cells, 2. antibody-dependent cellular cytotoxicity (ADCC), 3. HIV cure strategies, 4. broadly neutralizing antibodies, 5. Fc-optimized antibodies

## Abstract

The persistence of latent HIV-1 reservoirs remains the primary barrier to a cure. Shock and kill strategies aim to reactivate these reservoirs and eliminate them via effector cells, such as Natural Killer (NK) cells. However, chronic infection leaves NK cells exhausted. In this study, we investigated the interplay between cytokine-mediated NK cell activation and antibody-dependent cellular cytotoxicity (ADCC) across diverse HIV-1 subtypes. We demonstrated that while cytokine stimulation enhanced natural cytotoxicity, it simultaneously induces shedding of the Fc receptor CD16 via the metalloproteinase enzyme ADAM17. However, restoring CD16 expression through ADAM17 inhibition (TAPI-1) did not improve ADCC, suggesting that CD16 surface levels in NK cells is not the only limiting factor. On the other hand, Fc-engineered broadly neutralizing antibodies (bNAbs) with increased CD16 affinity, in particular LPLIL and GASDALIE, significantly enhance ADCC across the six different HIV-1 subtypes, regardless of NK cells activation, CD16 downregulation or limited surface Envelope expression in CD4T cells. These findings suggest that enhancing receptor affinity of bNAbs can bypass viral immune evasion and NK cell exhaustion, supporting their potential incorporation into HIV cure strategies.

**Importance:** Eradicating latent HIV reservoirs remains a global health priority, as it would liberate millions of people living with HIV (PLWH) from the economic and physiological burdens of lifelong antiretroviral therapy (ART). By utilizing a primary human cell model that resembles *in vivo* conditions across diverse viral strains, we identified that simply preventing CD16 receptor shedding from NK cells is insufficient to improve HIV-1 clearance. Instead, we demonstrated that enhancing the binding of antibodies to the CD16 receptor enables NK cells to overcome viral immune evasion and the loss of CD16 expression. This allows for potent elimination of infected cells through antibody-dependent cellular cytotoxicity (ADCC). These findings provide a clear strategy for designing more effective “kill” components in future therapeutic strategies.

## Introduction

Statistics from UNAIDS shows that Human Immunodeficiency Virus (HIV) continues to impact millions worldwide, with an estimated 40.8 million people living with the virus in 2024 (1–3). While antiretroviral therapy (ART) effectively suppresses viral replication and prevents the progression to acquired immunodeficiency syndrome (AIDS), it does not eliminate the virus, which persists in latent reservoirs (4–8).

Several strategies to deplete viral reservoirs are under investigation, such as the “shock and kill” strategy, which stimulates latently infected cells into active viral production using a latency-reversing agent (LRA) (9–11). This strategy aims to expose reactivated cells to immune-mediated clearance by cytotoxic lymphocytes, including CD8⁺ T cells and natural killer (NK) cells (9, 10, 12–14). Unfortunately, these effector cells often fail to eliminate reactivated cells, one reason being functional exhaustion, which is a consequence of chronic inflammation and prolonged immune activation (13, 15, 16). Current research is therefore focused on identifying strategies that can improve the cytotoxicity of effector lymphocytes, particularly NK cells which are a critical component of the innate immune system and essential for combating cancer and viral infections, including HIV (13, 17–20). NK cells eliminate HIV-infected cells through two primary mechanisms: natural cytotoxicity and antibody-dependent cellular cytotoxicity (ADCC). Natural cytotoxicity is initiated when NK cells recognize infected cells with reduced expression of major histocompatibility complex class I (MHC-I) molecules, a common viral immune evasion strategy (17, 21). ADCC, on the other hand, is mediated by the CD16 receptor (Fc-gamma receptor III - FcγRIIIa) on NK cells, which binds to the Fc region of antibodies binding HIV-infected cells (22). This interaction activates NK cells to release cytotoxic granules, resulting in the destruction of antibody-opsonized targets (23, 24).

To enhance the effectiveness of the “shock and kill” approach, broadly neutralizing antibodies (bNAbs) are being tested in combination with LRAs to facilitate the clearance of reactivated HIV-infected cells (25). Integrating bNAbs with LRAs has shown promise in preclinical and clinical studies, as it leverages the ability of bNAbs to target conserved epitopes of the HIV envelope glycoprotein trough their Fab portion opsonizing reactivated infected cells, and to promote ADCC through the Fc portion by engaging CD16 on NK cells and other immune effectors (22, 25, 26). However, during chronic HIV infection, persistent immune activation drives NK cell exhaustion, reducing cytotoxicity (13, 27). Additionally, the HIV accessory protein Vpu degrades Tetherin, allowing viral Env to detach from the cell surface. This decreases antibody binding limiting NK cell mediated ADCC (28). Adding to this challenge, CD16 is cleaved from the NK cell surface following activation by A Disintegrin and Metalloproteinase 17 (ADAM17) (29–34). To overcome this challenge, several groups have shown that enhancing the interaction of bNAbs with CD16 improves elimination of HIV-infected cells *in vitro* and in a mouse model (35, 36). In this work, we expand these studies using an HIV infection model with different HIV viral subtypes, primary CD4T cells and autologous NK cells to evaluate the impact of cytokine stimulation, CD16 expression in NK cells, HIV Env expression in CD4T cells, and affinity of bNABs to CD16 on ADCC.

## Results

### Cytokine-mediated NK cell activation reduces CD16 expression mediated by ADAM17

Cytokines such as interleukin-15 (IL-15), interleukin-12 (IL-12), and interleukin-18 (IL-18) play a central role in enhancing NK cell functionality. IL-15 is a key regulator of NK cell biology and cytotoxicity making it a key regulator of NK cell-mediated immunity in HIV infection (18, 37, 38). IL-12 is instrumental in NK cell maturation and the production of interferon-gamma (IFN-γ), a cytokine critical for antiviral responses (37). IL-18 synergizes with IL-12 and IL-15 to enhance NK cell proliferation, IFN-γ production, cytotoxicity and antiviral function (39–41).

We first evaluated the impact of cytokine stimulation on ADCC. NK activation, measured as CD69 expression and IFN-γ production, was assessed after treatment of NK cells with IL-15 alone or in combination with IL-12 and IL-18. As expected, cytokine stimulation led to NK cell activation (Fig. 1A,B) and IFN-γ production (Fig. S1A,B), with the highest induction mediated by IL-12/IL-15/IL-18 treatment. Triple cytokine treatment was associated with a small albeit significant reduction on viability of NK cells (Fig. 1C). Interestingly, cytokine stimulation led to a reduction on the surface expression of CD16 (Fig. 1D and Fig. S1D). Surface CD16 expression was negatively associated with CD69 and IFN-γ induction (Fig. 1E and Fig. S1E). NK activation and loss of CD16 expression was independent on biological sex or age (Fig. S1F-K).

**Fig 1.**
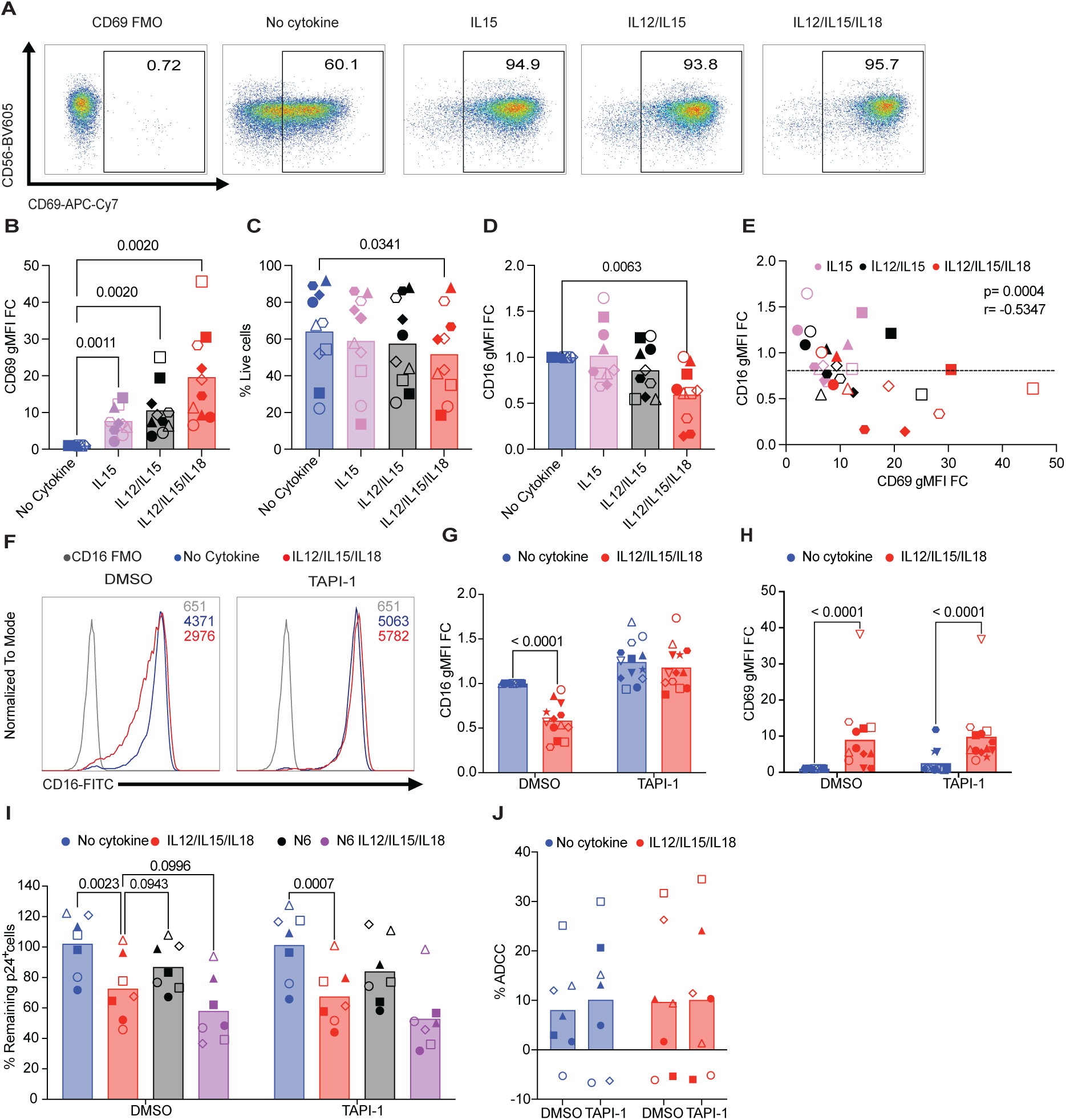
Cytokine stimulation enhances NK cell activation while downregulating CD16 expression through ADAM17-dependent shedding. (**A**) Representative flow plots of CD69 induction of NK cells stimulated with IL-15, IL-12/IL-15, or IL-12/IL-15/IL-18. (**B**) CD69 induction across donors (n=10). (**C**) Viability upon different treatments. (**D**) Changes in CD16 expression across multiple donors upon cytokine treatment. (**E**) Spearman correlation analysis between CD16 expression and CD69 induction. (**F**) Representative histogram of CD16 expression upon triple cytokine treatment in the presence of TAPI-I (right panel) or DMSO control (left panel). (**G**) CD16 expression in donors (n=13) stimulated with cytokines in the absence or presence of TAPI-1. (**H**) Effects of TAPI-I on NK cell activation. (**I**) Percentage remaining p24^+^ cells upon coculture of HIV-infected CD4^+^T cells with autologous NK cells from donors (n=7) pre-stimulated with cytokines in the presence of TAPI-1 or DMSO control. (**J**) Percentage ADCC calculated as indicated in methods from data shown in (**I**). Statistical significance was calculated using one-way ANOVA with multiple comparison compared to the no cytokine condition for panel (**B-D**), two-way ANOVA with multiple comparison compared to the IL12/IL15/IL18 condition for panel (**I**), and two-way ANOVA for panel (**J**). Open symbols represent female donors and closed symbols male donors.

CD16 downregulation following cytokine activation is a known regulatory mechanism mediated by ADAM17-cleavage (32). However, its impact on ADCC, distinct from natural cytotoxicity, remains poorly defined in primary HIV models. We first confirmed that the metalloproteinase inhibitor TAPI-I restored CD16 expression upon IL12/IL15/IL18 stimulation (Fig. 1F, G) without altering NK activation (Fig. 1H). We next evaluated whether restoring CD16 expression was associated with increased ADCC of HIV-infected autologous CD4T cells (Fig. S2A). Briefly, primary CD4T cells infected with the HIV strain NL4-3 (subtype B) were co-cultured with autologous NK cells and natural cytotoxicity and ADCC were measured as a reduction in the percentage of cells expressing HIV p24Gag and downregulating CD4 by flow cytometry (Fig. S2B) (42). IL12/IL15/IL18 stimulation enhanced natural cytotoxicity and TAPI-1 did not influence this killing mechanism (Fig. 1I and Fig. S2C, D). As expected, the presence of the CD4 binding site bNAb N6 promoted ADCC. Interestingly, IL12/IL15/IL18 activation or the TAPI-1 inhibitor did not enhance ADCC (Fig. 1J, Fig. S2E, F). These results indicate that while cytokine stimulation enhances natural cytotoxicity, it does not influence ADCC. Furthermore, restoring surface CD16 expression by inhibiting ADAM17 does not improve ADCC, even in the context of IL12/IL15/IL18-activated NK cells, highlighting that restoring the CD16 receptor density on NK cells does not linearly translate to improved ADCC.

### Enhancing the affinity of bNAbs to CD16 enhances ADCC

Given that restoring surface CD16 expression alone is not sufficient to enhance ADCC, we evaluated whether enhancing antibody affinity for the CD16 receptor could restore ADCC even in the context of reduced surface CD16 expression. To test this, we used the CD4-binding site bNAb VRC07-523-LS (WT) along with a panel of antibodies with mutations on the Fc region that either enhances binding to CD16 including VRC07-AAA, VRC07-LPLIL, and VRC07-GASDALIE; or a mutant with reduced CD16 affinity VRC07-GRLR (21, 35, 43, 44) (Fig. 2A). Using the Jurkat-Lucia NFAT-CD16 reporter cell line, we confirmed that FcR enhancing mutations enhance CD16 binding relative to wildtype while the negative control GRLR lacks activity (Fig. 2B). Next, we assessed the ability of each construct to promote NK cell activation through its interaction with CD16 independent of antigen recognition. To that end, non-tissue culture-treated plates were pre-coated overnight with each of the antibodies. The next day, isolated NK cells were plated overnight, and NK activation was measured by analyzing expression of the activation marker CD69 and cytokine secretion. All antibodies except GRLR were able to induce NK activation, and when adjusted for multiple comparison, all FcR enhancing mutations significantly induce NK activation (Fig. 2C). Some minor toxicity was observed for LPLIL and GASDALIE constructs (Fig. 2D). Next, we evaluated the ability of each construct to promote cytokine secretion using an ultrasensitive 10-plex cytokine array measuring IFN-γ, TNF-α, IL-1β, IL-4, IL-5, IL-6, IL-8, IL-10, IL-12p70, and IL-22 (Fig. 2E-O).

**Fig 2.**
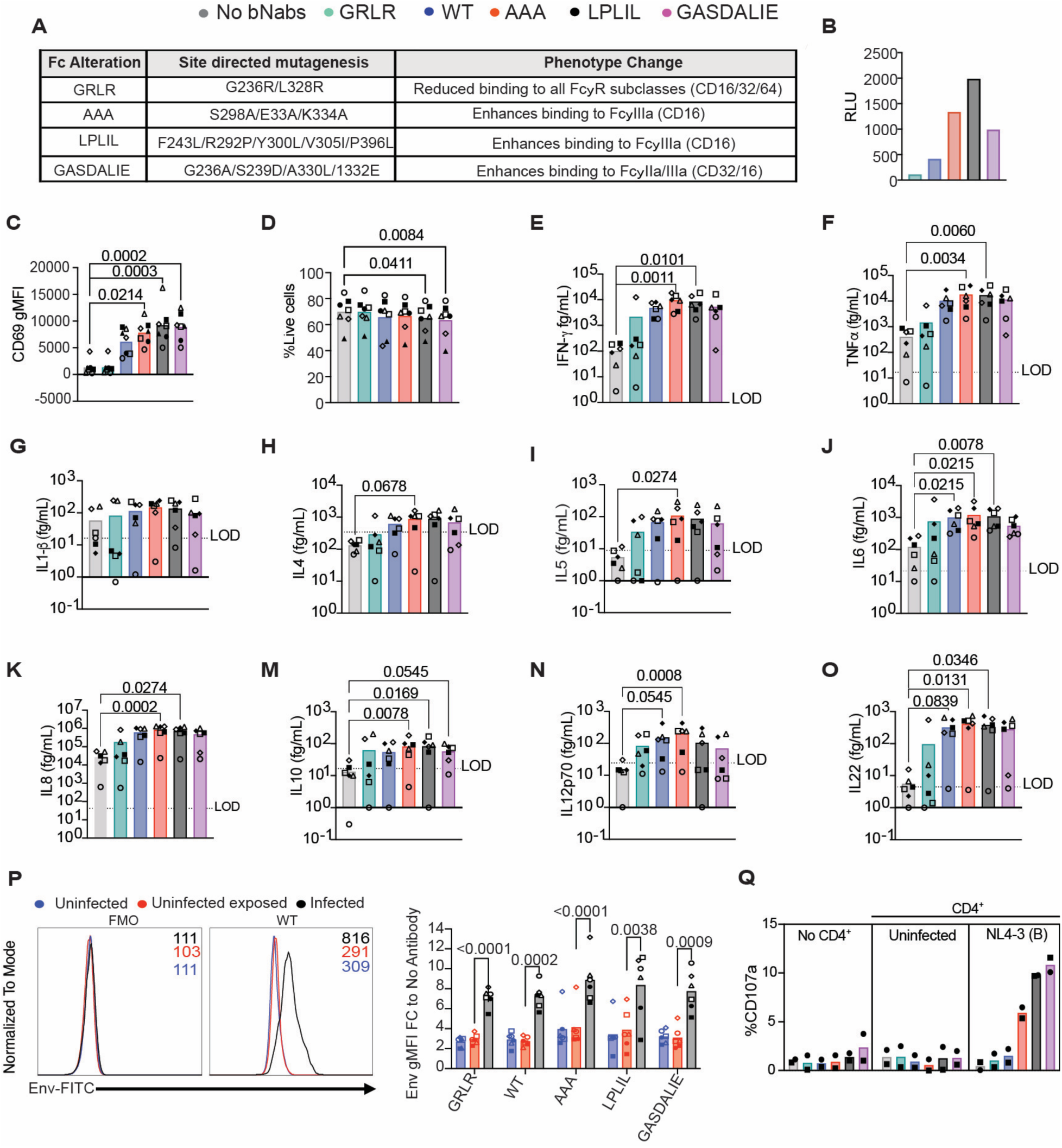
Functional activity, cytokine secretion, and Fc binding of VRC07-523-LS and Fc variants. (**A**) Table summarizing VRC07-523LS antibody mutants with optimized Fc biding including specific mutations and effects on CD16 binding. (**B**) CD16 reporter activity in Jurkat-Lucia NFAT-CD16 cells exposed to antibody variants. (**C**) CD69 induction across donors (n=7). (**D**) Viability upon different conditions. (**E-O**) Supernatants from NK cells cultured overnight with WT VRC07-523-LS or Fc mutant antibodies were analyzed for cytokine secretion (fg/mL). Levels of (**E**) IFN-γ (Limit of detection (LOD) 1fg/mL), (**F**) TNF-α (LOD 16.7fg/mL), (**G**) IL-1β (LOD 16.3fg/mL), (**H**) IL-4 (LOD 339.1fg/mL), (**I**) IL-5 (LOD 8.7fg/ml), (**J**) IL-6 (LOD 21.1fg/mL), (**K**) IL-8 (LOD 41.7fg/mL), (**M**) IL-10 (LOD 16.8fg/mL), (**N**) IL-12p70 (LOD 24.7 fg/mL), (**O**) IL-22 (LOD 4.5 fg/mL) (n=6). (**P**) Representative flow cytometry histograms showing VRC07-523LS binding to surface Env on uninfected (blue), uninfected exposed (red), or HIV-infected cells (black). A FMO control is included (grey). Quantification of Env binding for the different antibodies (n=7). (**Q**) Surface expression of CD107a upon co-culture of NK cells alone (no CD4^+^), or NK cells co-cultured with uninfected autologous CD4^+^ or NL4-3 infected CD4^+^ T cells in the presence of the indicated antibody (n=2). Statistical significance was assessed using a Friedman test with multiple comparisons compared to unstimulated condition for panels (**C-O**), and two-way ANOVA with multiple comparisons for panel (**P**). Open symbols represent female and closed symbols male donors.

When adjusted for multiple comparison, both the AAA and LPLIL mutants induced significantly higher secretion of several cytokines including IFN-γ (Fig. 2E), TNF-α (Fig. 2F), IL-6 (Fig. 2J), IL-8 (Fig. 2K), IL-10 (Fig. 2M), and IL-22 (Fig. 2O). Furthermore, the AAA mutant induced higher levels of IL-4 (Fig. 2H), IL-5 (Fig. 2I), and IL-12 (Fig. 2N). None of the constructs induced IL-1β (Fig. 2G). Notably, some cytokine secretion was also detected with the GRLR mutant in certain donors, despite its design to abrogate CD16 binding. This residual activity likely reflects its remaining receptor engagement capacity, suggesting that even minimal Fc-receptor interaction can trigger detectable cytokine responses. We next evaluated whether changes in the Fc region interfere with binding of the Fab region of the bNAbs to HIV Env in the surface of HIV-infected primary CD4T cells. Using flow cytometry, we validated that VRC07-523-LS binds to the surface of HIV-infected CD4T cells but not to uninfected cells or uninfected exposed cells in the same culture. Importantly, none of the Fc mutations altered Fab binding to surface HIV Env (Fig. 2P). Next, we evaluated whether antigen is required to promote NK degranulation with these constructs. To that end, NK cells were cultured alone or in the presence of autologous uninfected CD4T cells or HIV-infected CD4T cells in combination with each of the bNAbs. 4-hours later, NK degranulation was measured by the induction of the degranulation marker CD107a. We confirmed that NK cells only degranulate in the presence of HIV-infected cells with increased degranulation observed with bNAbs with enhance CD16 affinity (Fig. 2Q)

Finally, we evaluated the ability of each construct to mediate ADCC on HIV-infected primary CD4T cells. Compared to WT bNAb, all three constructs with enhanced Fc binding increased killing of HIV-infected cells regardless of IL12/IL15/IL18 treatment while the GRLR mutant lacked activity (Fig. 3A and Fig. S3A-B, S4A-F). As observed with N6, restoring CD16 expression with TAPI-I treatment did not significantly enhanced killing (Fig. 3B and Fig. S4G-L). Hierarchical clustering based on the % of remaining HIV-infected CD4T cells demonstrated that treatments clustered by either IL12/IL15/IL18 treatment (Cluster 1 vs Cluster 2), or antibody affinity to CD16 (Cluster 3 vs Cluster 4), with ADAM17 inhibition having no effect on NK-killing of HIV-infected CD4T cells (Figure 3C and Supplemental Figure 5). We specifically evaluated ADCC as indicated in the Methods section. Compared to WT, all the constructs with enhanced Fc binding increased ADCC regardless of cytokine treatment (Figure 3D) or TAPI-I treatment (Figure 3E) while GRLR showed no ADCC. Hierarchical clustering demonstrated that ADCC clustered primarily by antibody affinity to CD16 (Cluster 3 vs Cluster 4) with IL12/IL15/IL18 treatment or ADAM17 inhibition having minimal effect on ADCC (Fig. 3F). Finally, we used the Bliss independence model to evaluate potential synergy of each bNAb with IL12/IL15/IL18 treatment in the presence or absence of TAPI-1. This model uses probability to determine if two treatments are acting through synergistic mechanisms. Values above 0 are considered synergistic, while values below 0 are antagonistic and those equal to 0 are consider independent or additive (45). Regardless of TAPI-I treatment, bNAbs and IL12/IL15/IL18 stimulation act independently (Fig. S6A-B). In conclusion, these findings highlight that while enhancing surface CD16 expression alone does not improve ADCC, increasing bNAb affinity for CD16 significantly enhances ADCC activity regardless of the activation status of NK cells.

**Fig. 3.**
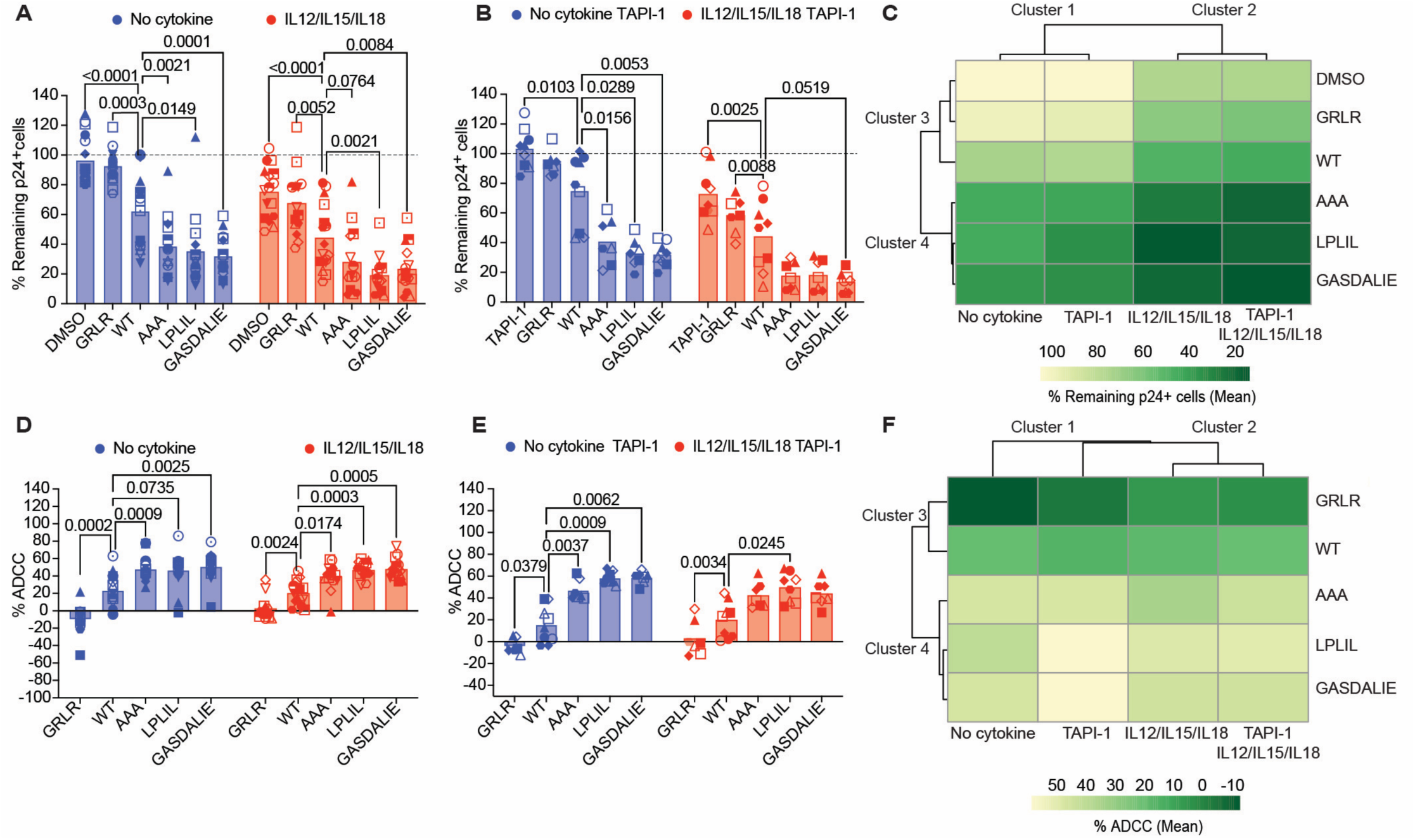
Enhancing antibody affinity to CD16 improves NK cell-mediated ADCC against HIV-infected cells independently of cytokine activation or ADAM17 inhibition. Percent of remaining HIV p24⁺ infected CD4⁺ T cells following co-culture with autologous NK cells from donors pre-treated with or without cytokines (IL-12/IL-15/IL-18) (n=16) (**A**) and ADAM17 inhibitor TAPI-1 (n=7) (**B**), in the presence of the indicated antibody. (**C**) Hierarchical cluster analysis of percentage of remaining p24⁺ cells in different conditions shown in A-B. ADCC activity under the same treatment conditions in the absence (**D**) or the presence (**E**) of the ADAM17 inhibitor TAPI-1. Statistical significance was determined using two-way ANOVA with multiple comparison compared to WT. Open symbols represent female and closed symbols male donors. (**F**) Hierarchical cluster analysis of ADCC in different conditions shown in **D-E**. Clusters were done using ClustVis.

### Enhancing Fc/CD16 interaction improves killing even with limited binding to surface envelope

Given the extensive genetic diversity of the HIV-1 *env* gene, we evaluated VRC07-523LS and its Fc mutants against five viral subtypes. While Fc engineering has shown to enhance ADCC (35, 36), its efficacy has not been systematically characterized across a diverse genetic landscape of HIV-1 subtypes or with treatments that promote CD16 downregulation. All viruses selected were sensitive to neutralization by VRC07-523LS, albeit with varying half inhibitory concentration (IC50) ranging from 0.008 µg/mL to 0.419 µg/mL (Fig. 4a, Fig. S7A). Likewise, surface Env binding varies across viral subtypes with Fc mutations not affecting binding (Fig. 4B, Fig. S7B–F). As previously shown, surface binding to HIV-infected cells correlated with neutralization potency, both IC50 and IC80, although statistical significance was limited due to donor-to-donor variability (46) (Fig. 4C, Fig. S7A).

**Fig. 4.**
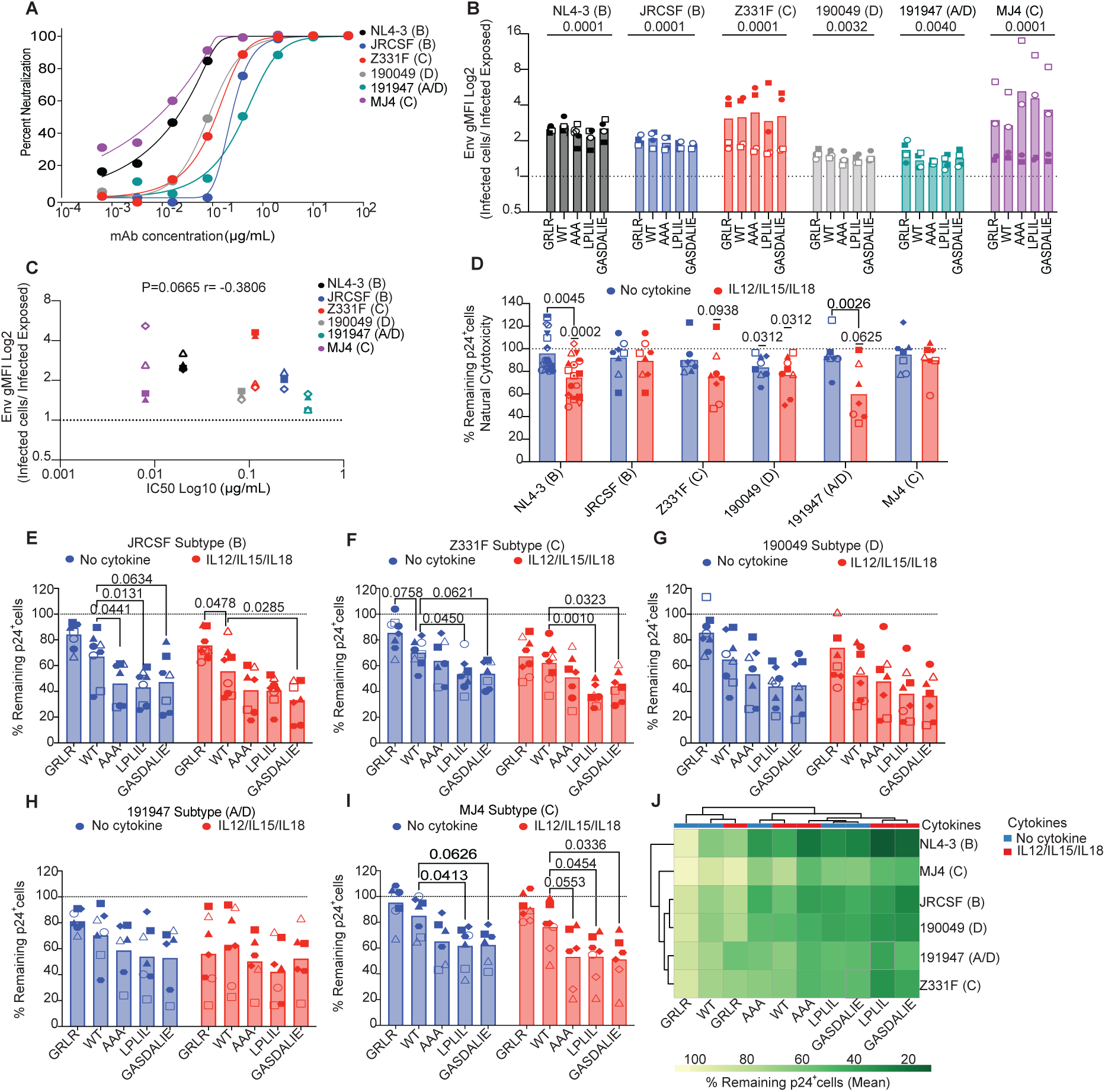
Effects of Fc-optimized VRC07-523LS mutants across HIV-1 subtypes. (**A**) Neutralization curves of VRC07-523LS against indicated HIV-1 viral strain. (**B**) Env binding of the different antibodies to HIV-infected CD4T cells with the indicated viral strain (n=3-4). (**C**) Spearman correlation between Env binding and neutralization IC_50_ values. (**D**) Percentage of remaining p24^+^ cells following natural cytotoxicity by autologous NK cells pretreated with a combination of cytokines or no cytokine control (n= 6-16). (**E-I**) Percentage of remaining p24+ cells for each of the indicated viruses in the presence of the different antibodies (n= 7-8). Statistical significance was assessed using a one-sample t test with a hypothetical value set to 1 for panel (**B**), wilcoxon signed rank test with hypothetical value set to 100 for panel (**E**), <u>and two</u>-way ANOVA with multiple comparison compared to WT for panels (**E–I**). Open symbols represent female and closed symbols male donors. (**J**) Hierarchical cluster analysis of the percentage remaining p24^+^ cells. Clusters were done using ClustVis.

Next, we evaluated natural cytotoxicity of NK cells against the different viral strains and the effect of IL12/IL15/IL18 stimulation. IL12/IL15/IL18 stimulation significantly enhanced natural cytotoxicity relative to untreated NK cells only for NL43 (B) and 191947 (A/D) (Fig. 4D). As expected, the ability of NK cells to kill HIV-infected cells upon cytokine treatment was highly correlated with the ability of the virus to downregulate MHC-I from the surface (Fig. S8). Across the multiple viral subtypes, Fc-optimized variants mediated stronger killing of HIV-infected CD4T cells compared to WT, whether NK cells were activated or not with IL12/IL15/IL18 (Fig. 4E–I). Hierarchical clustering analysis across the six viral subtypes revealed that Fc-optimized variants, particularly LPLIL and GASDALIE, consistently demonstrated the broadest and most potent activity against diverse HIV-1 strains (Fig. 4J, Fig. S9).

We next compared ADCC activity of VRC07-523LS WT and mutants across the six HIV-1 strains in the presence or absence of IL12/IL15/IL18. Similar to NL4-3, VRC07-523LS mediated measurable ADCC against all isolates and cytokine stimulation did not influence ADCC except for 191947 (A/D) (Fig. 5A). This viral strain is particularly sensitive to cytokine-mediated natural cytotoxicity (Fig. 4D) and the addition of bNAbs did not seem to enhance killing (Fig. 4H). Upon IL12/IL15/IL18 stimulation, there was a significant positive correlation between ADCC and envelope binding (Fig. 5B) but not with unstimulated NK cells. These data suggest that activated NK cells rely on surface envelope expression to mediate ADCC, potentially due to limited CD16 expression (Fig. 1). The Fc-optimized bNAbs consistently mediated higher ADCC activity across all tested viruses (Fig. 6A–E). Hierarchical clustering demonstrated that all Fc-optimized variants grouped closely together, and distinct clusters were observed under IL12/IL15/IL18-pretreated versus untreated conditions (Fig. 6F). As for NL4-3, synergy analysis using the Bliss independence model showed an additive effect between IL12/IL15/IL18 stimulation and bNAb activity across multiple viral subtypes (Fig. S10).

**Fig. 5.**
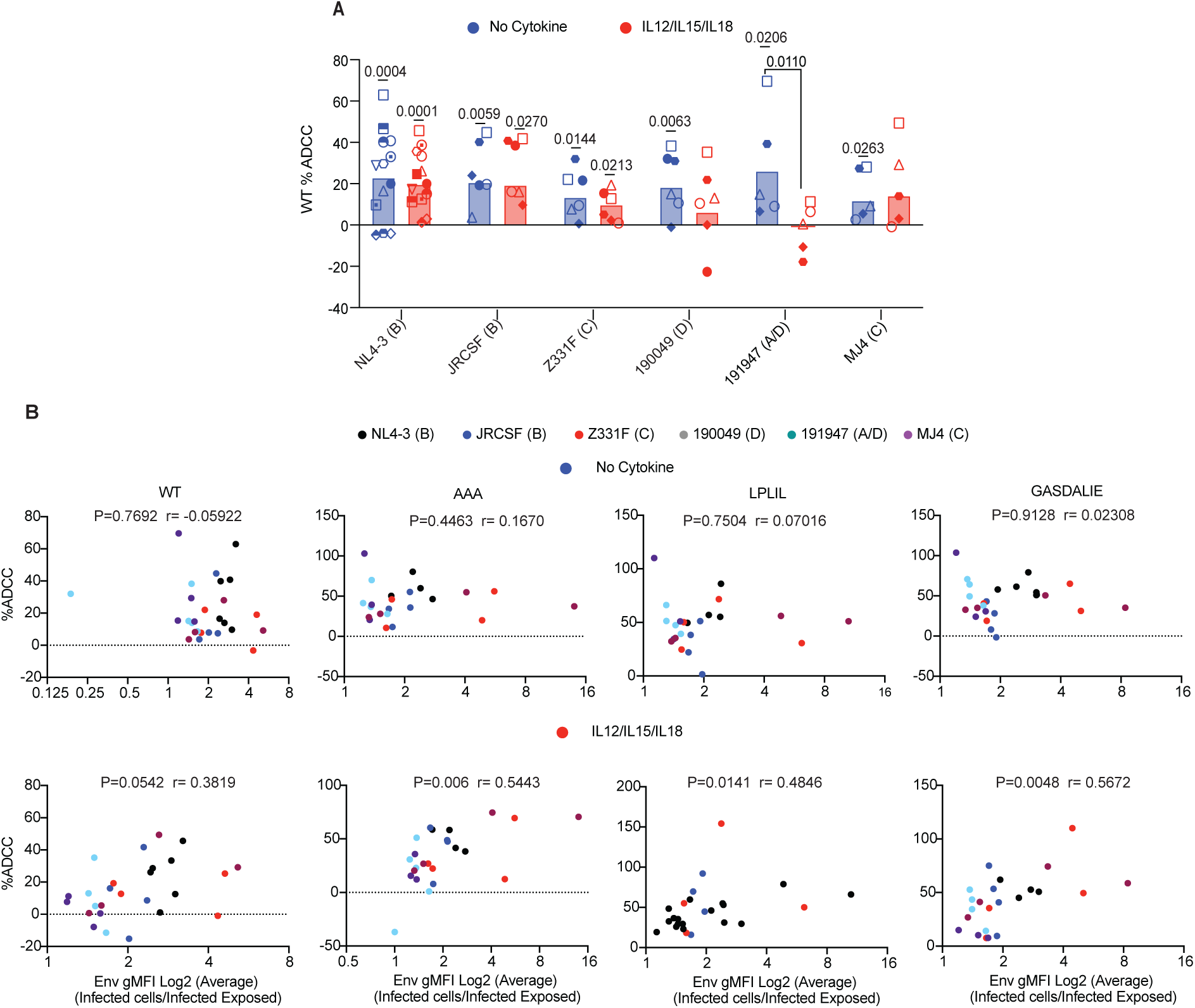
Cytokine activation of NK cells requires higher levels of surface Env binding to promote ADCC. (**A**) Percentage of ADCC mediated by VRC07-523LS for each of the viral strains upon pretreatment of NK cells with the combination of cytokines or no cytokine control (n=6-16). Statistical significance between no cytokine and IL12/IL15/IL18 was determined by two-way ANOVA and Wilcoxon signed rank test with hypothetical value set to 0 for each condition. Open symbols represent female and closed symbols male donors. (**B**) Spearman correlation between the percentage of ADCC and surface Env binding for the indicated antibodies and NK cells pretreated with no cytokine (top) or IL12/IL15/IL18 (bottom). Each virus is indicated with a different color (n=4-6).

**Fig 6.**
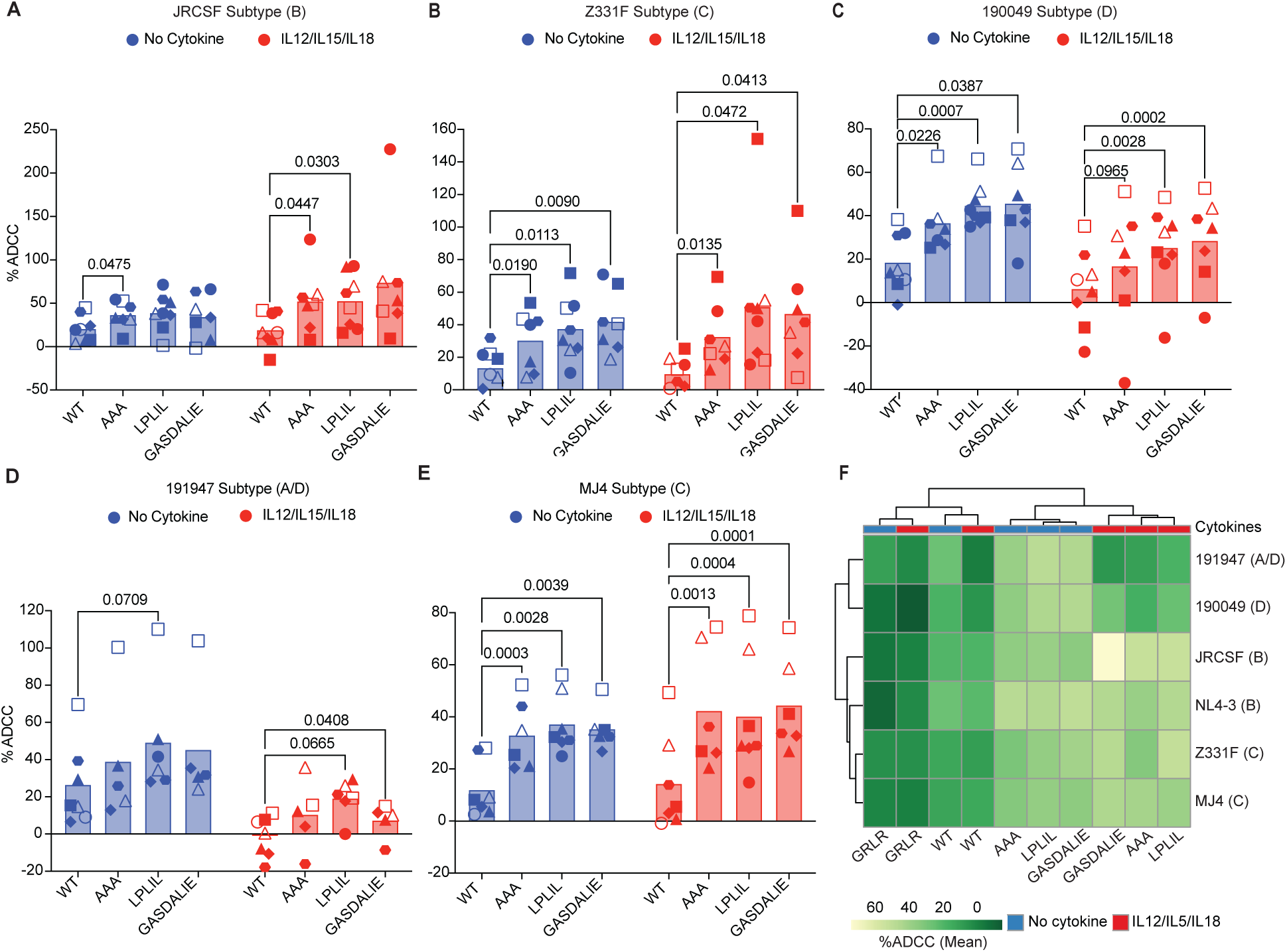
Increased ADCC mediated by the Fc-optimized VRC07-523LS mutants across diverse HIV-1 subtypes. (**A-E**) Percentage of ADCC mediated by VR07-523LS and Fc-optimized variants across the indicated viral subtypes (n=7-8). Statistical significance was determined two-way ANOVA with multiple comparison compared to WT. Open symbols represent female and closed symbols male donors. (**F**) Hierarchical cluster analysis of ADCC. Clusters were done using ClustVis.

## Discussion

In this study, we examined how cytokine stimulation, CD16 receptor modulation, and Fc-optimized bNAbs influence NK cell-mediated killing across diverse HIV-1 subtypes. We further expanded previous research by evaluating these strategies across diverse HIV-1 subtypes and using an autologous HIV-infection model. While prior studies observed that IL-15 increase NK cell-mediated cytotoxicity against HIV-infected CD4T cells, they utilized readouts that capture combined cytotoxic effects (natural cytotoxicity and ADCC) and did not exclusively evaluate ADCC (47). Similarly, recent evaluations of Fc-optimized bNAbs have used PBMCs rather than purified NK cells, making it difficult to isolate FcγRIIIa-mediated ADCC from other immune mechanisms (36). Furthermore, both studies utilized transformed cell lines as targets, which are less resistant to lysis than HIV-infected primary CD4T cells. Overall, while bNAbs may provide broad neutralization of viral strains, our results demonstrate that Fc-affinity is the primary driver of ADCC potency, with variants like LPLIL and GASDALIE maximizing clearance across diverse HIV-1 strains. Crucially, this enhanced activity remains strictly antigen-dependent, augmenting degranulation against HIV-infected cells without triggering non-specific responses.

Our findings reveal that while IL12/IL15/IL18 stimulation robustly activates NK cells, as indicated by induction of CD69 and IFN-γ production, it is not sufficient to enhance killing of HIV-infected cells to all the viral strains analyzed nor improves ADCC. Natural cytotoxicity activity upon IL12/IL15/IL18 stimulation is highly correlated with the ability of the viral strain to downregulate MHC-I from the cell surface of HIV-infected CD4T cells (Fig.S8C) (22). On the other hand, IL12/IL15/IL18 stimulation mediates downregulation of the Fc receptor CD16 critical for ADCC. An ADAM17 inhibitor, which successfully restored CD16 surface expression, did not enhance ADCC (Fig. 1F-J). Upon IL12/IL15/IL18 stimulation, NK cells rely primarily in binding to surface envelope to induce ADCC (Fig. 5B). Reduced CD16 expression likely alters receptor dynamic and prevents saturation under our experimental conditions. On the other hand, the higher levels of CD16 in non-stimulated NK cells may exhibit saturated CD16 occupancy, maximizing antibody engagement and signaling (Fig. 5B). This saturation could overcome limited binding to surface envelop expression observed in some of the viral strains. Furthermore, we found that IL12/IL15/IL18 stimulation and bNAb treatment had an additive rather than a synergistic effect. This indicates that natural cytotoxicity and ADCC utilize independent but converging killing mechanisms. Therefore, the benefit of one pathway is not exponentially increased by the other. While Fc engineering is a known strategy to enhance antibody function, its efficacy across the vast genetic landscape of HIV-1 subtypes particularly when NK cells are in a state of lower CD16 expression has not been systematically characterized (35).

From the Fc modifications tested, GASDALIE and LPLIL consistently demonstrated superior cytolytic activity over AAA across all viral strains, independent of cytokine priming. The GASDALIE (G236A/S239D/A330L/I332E) promotes stabilization of the Fc-FcγRIIIa interface which results in increasing receptor affinity and enhance NK cells activation (35, 44, 48). Compared to wildtype, GASDALIE has a 431-fold increase in affinity to FcγRIIIa_F158_ (48). The LPLIL (F243L/R292P/Y300L/P396L) has been shown to improve the binding to the CD16A receptor by improving the affinity to the high affinity Val158 and low affinity Phe158 in CD16 while keeping the same binding to CD32B, an inhibitory receptor (43). These mutations modify the lower hinge and CH2 domains to improve FCγR engagement while preserving Fc stability and antigen recognition (21, 43). In contrast AAA (S298A/E333A/K334A) provides a minor increase to the CD16a affinity but does not stabilize the Fc-FCγR interface. (44). The lack of AAA to stabilize the Fc-FCγR interface may result in lower activity and may explain our results indicating that GASDALIE and LPLIL outperform AAA across the various HIV-1 subtypes. Notably, despite improved cytotoxic function, we did not observe a significant difference in cytokine secretion between WT and Fc-optimized antibodies. This might suggest that Fc optimization augment cytolytic activity without broadly amplifying inflammatory cytokine responses, potentially minimizing off-target effects.

Our study has some caveats. While our study utilizes an *in vitro* super-infection model to allow for high throughput testing across the viral subtypes, it specifically enabled us to characterize the impact of cytokine-induced CD16 downregulation on ADCC, a critical barrier that mirrors the receptor loss seen in chronic NK cell exhaustion (13, 27). Future studies using LRAs and cells from ART suppressed PLWH must confirm if these Fc-optimized bNAbs can overcome latent reservoir challenges. This includes not only low Env density but also the chronic NK cell exhaustion prevalent in PLWH, where persistent CD16 shedding and internal signaling defects may reduce the efficiency of antibody-mediated clearance compared to what we observed in our primary cell model. Additionally, we acknowledge that using NK cells isolated directly from cryopreserved PBMCs may impact activation kinetics compared to fresh cells. However, to minimize variability, we utilized autologous systems and strictly standardized isolation and cytokine pretreatment protocols. The robust natural cytotoxicity and ADCC observed across multiple viral subtypes demonstrate that the NK cells remained functionally competent and responsive within our assay system. While our *in vitro* model demonstrated that the Fc-optimized bNAbs amplify NK mediated ADCC against the six viral subtypes, studies in viremic, ART naïve macaques show no therapeutic advantage for Fc-optimized bNAbs (49, 50). This discrepancy can be attributed by the uncontrolled viremia that favors neutralization or antibody mediated binding of the free virus, while disfavoring ADCC and antibody-dependent cellular phagocytosis (ADCP). Mutations enhancing Fc-CD16 interaction may have a more favorable profile in cure strategies in the presence of ART, which antibody effector functions opsonizing reservoir cells may promote better elimination.

Collectively, our results support the potential of Fc-optimized bNAbs to enhance NK cell-mediated ADCC against HIV-infected cells. Their consistent efficacy across diverse viral subtypes and independence from cytokine priming make them attractive candidates for inclusion in cure strategies. Simultaneously, leveraging cytokines offer a parallel axis for enhancing NK cell activation. For example, the IL-15 superagonist N-803 is currently undergoing clinical trials in ART-suppressed PWH and has been shown to enhance NK cell cytotoxicity reinvigorating exhausted NK cells (51, 52). In conclusion, our findings highlight the potential of high-affinity Fc-engineering to overcome biological barriers in a vast genetic landscape and compensate for the CD16 receptor loss seen in activated NK cells to develop multi-pronged strategies for HIV control. Translational validation in ART-suppressed individuals will be required to confirm these effects within the persistent reservoir.

## Material and Methods

### Reagents

Recombinant human IL-2 (rhIL-2) and rhIL-15 were provided by the BRB/NCI Preclinical Repository. Recombinant human IL-12 (rhIL-12) was obtained from PeproTech, and rhIL-18 from R&D Systems. The following reagents were acquired through the NIH HIV Reagent Program, Division of AIDS, NIAID, NIH: Nelfinavir, Raltegravir, and the HIV-1 Strain NL4-3 Infectious Molecular Clone (pNL4-3), ARP-114, and TZM-bl cells. Additional HIV-1 infectious molecular clones included strain JR-CSF (pYK-JRCSF), ARP-2708, contributed by Dr. Irvin S. Y. Chen and Dr. Yoshio Koyanagi; Human Immunodeficiency Virus Type 1 Z331F Infectious Molecular Clone (SGA 11), ARP-13249, contributed by Dr. Eric Hunter; the HIV-1 MJ4 infectious molecular clone (pMJ4) was gifted by Thumbi Ndung’u, Boris Renjifo, and Max Essex (cat# 6439) (53). The HIV-1 primary isolates p191947 and p190049 were a generous gift from Dr. Beatrice H. Hahn. The monoclonal antibody VRC07-523LS was provided by the Vaccine Research Center (VRC), NIH. Additional reagents included Dynabeads Human T-Activator αCD3/CD28 (cat# 11131D) were purchased from ThermoFisher, YU2 gp120 protein (ImmuneTech), B3T buffer, 1% Tween 20 wash buffer, QUANTILuc luciferase, SpectraMax Glo Steady-Luc reporter assay reagent (Molecular Devices), DEAE-dextran (Sigma-Adrich), indinavir (Sigma-Aldrich), and antibiotic-free IMDM and DMEM media.

#### Generation of Infectious HIV-1 Molecular Clones

Infectious HIV-1 virus was generated using HEK293FT. Cells were maintained in DMEM (high-glucose, without L-glutamine or sodium pyruvate) and transfected using the calcium phosphate (CaPO_4_) method, as previously described (54, 55). Briefly, plasmid DNA was mixed with a CaPO_4_ solution, incubated to allow precipitate formation, and then added dropwise to HEK293FT cells. After 48 hours, the supernatants, containing the virus, were collected, filtered through a 0.45 µm membrane to remove cellular debris, and stored at -80. Viral titers were determined by infecting day 7-activated CD4⁺ T cells and performing titration assays.

#### PBMCs isolation

Peripheral blood mononuclear cells (PBMCs) were isolated from buffy coats obtained from HIV-negative donors through the Gulf Coast Regional Blood Center. PBMCs were separated using a density gradient centrifugation method with Lymphoprep (STEMCELL Technologies, cat# 07851). The cells were then washed three times with PBS containing 2mM EDTA to remove residual contaminants. Finally, PBMCs were resuspended in complete media, consisting of RPMI 1640 supplemented with 10% fetal bovine serum (FBS, Gibco), 1% L-glutamine, and 1% Penicillin/Streptomycin (Gibco).

#### Binding profiles of Fc enhancing and abrogating mutations to VRC07-523 LS

ELISA plates were coated with 2 μg/mL YU2 gp120 protein (ImmuneTech) in PBS and incubated overnight at 4°C. After washing 6x with 1% Tween 20 wash buffer, plates were blocked using 200 μL B3T buffer for 1 hour at 37°C. Plates were then washed again and 100 μL antibodies were added to wells after being serially diluted in B3T buffer starting at 10 μg/mL. After a 1-hour incubation at 37°C, cells were resuspended in antibiotic free IMDM media at 1×10^6^ cells/mL and 100 μL was added to each well. Plates were incubated overnight at 37°C before being read using QUANTILuc luciferase.

#### In vitro NK activation using bNABs

Non-tissue culture-treated 24 well plates (Falcon, A Corning Brand), were pre-coated overnight at 37°C in a 5% CO_2_ with 10μg/mL of VRC07-523-LS and mutants in PBS. Next, day the NK cells were added 500,000 cells per well and incubated overnight at 37°C in a 5% CO_2._ The next day, cells were washed in PBS and incubated with Human Fc Block (BD Biosciences, cat#564220) for 10 minutes at room temperature. Cells were then stained with surface markers, including anti-human CD3 PerCP/Cy5.5 (BD Biosciences, cat# 362506), anti-CD16 FITC (BD Biosciences clone 3G8, cat# 555406), anti-human CD56 BV605 (BioLegend, cat# 318333), anti-human CD69 APC-Cy7 (BioLegend, cat# 310914), and eBioscience™ Fixable Viability Dye eFluor™ 450 (Thermo Fisher Scientific, cat# 65-0863-18) in FACS buffer (PBS with 2% FBS). Staining was conducted for 30 minutes at 4°C. Supernatants from the same cultures were collected for cytokine analysis as described below.

#### Cytokine analysis

Frozen supernatants were thawed, and the assay was performed according to Quanterix Human 10-Plex protocol. Briefly, samples were diluted 1:4 with Sample Diluent provided with the kit incubated on the 96-well pre-spotted plate for 2 hours at RT and shaking at 525 rpm. Then plate was washed with Quanterix wash buffer (cat# 1863332) using a plate washing and then incubated first with biotinylated antibody for 30 minutes, washed and then incubated with streptavidin-HRP for 30 minutes each at RT and shaking at 525 rpm. Next the plate was washed with a 5-step wash, then the SuperSignal provided from the kit was added, and plate was read. Ten cytokines were measured using Quanterix SP-X Corplex Cytokine Panel (IFN-γ, IL-1β, IL-4, IL-5, IL-6, IL-8, IL-10, IL-12p70, IL-22, TNF-α) (cat# 85-0329).

#### TAPI-1 preparation

TAPI-1 inhibitor (Selleck Chemicals, cat# S7434) was initially dissolved at 100 mM in DMSO. An intermediate dilution was prepared in complete media at a concentration of 1 mM. The cells were treated with the final concentration of TAPI-1 inhibitor at 10 μM, with DMSO maintaining a final concentration of 0.01%.

#### NK cells Phenotype

NK cells were isolated from PBMCs using the EasySep™ Human NK Cell Isolation Kit (STEMCELL Technologies, cat# 17955). NK cells were treated with 50 ng/ml IL-15, 10 ng/mL IL-12 and/or 50 ng/mL IL-18 as indicated in the presence or absence of 10 μM TAPI-I or DMSO control. For IFN-γ, NK cells were incubated for 1hr and then protein transporter inhibitor cocktail at 1x (eBioscience Protein transport inhibitor cocktail (500x), cat# 00-4980-03) was added. After 24 hours of incubation at 37°C, cells were stained for surface and intracellular markers.

Cells were washed in PBS and incubated with Human Fc Block (BD Biosciences, cat# 564220) for 10 minutes at room temperature. Cells were then stained with surface markers, including anti-human CD3 BV786 (BD Biosciences, cat# 563918), anti-CD16 FITC (BD Biosciences, clone 3G8, cat# 555406), anti-human CD56 PerCP/Cy5.5 (BioLegend, cat# 362506), anti-human CD69 APC-Cy7 (BioLegend, cat# 310914), and eBioscience™ Fixable Viability Dye eFluor™ 450 (Thermo Fisher Scientific, cat# 65-0863-18) in FACS buffer (PBS with 2% FBS). Staining was conducted for 30 minutes at 4°C. For IFN-γ, surface staining anti-CD3-BV786 (BD Biosciences, clone SP34-2, cat# 563800), anti-CD16 FITC (BD Biosciences, clone 3G8, cat# 555406), and anti-CD56-BV605 (Biolegend, clone HCD56, cat# 318334). Following surface staining, cells were washed, fixed, and permeabilized using Cytofix/Cytoperm™ (BD Biosciences, cat# 554722) according to the manufacturer’s protocol. Intracellular staining was then performed using anti-IFN-γ APC-A (BD Biosciences, clone OKT4, cat# 17-0048-42). After staining, cells were washed and resuspended in 3% PFA. Samples were acquired using a BD LSR Fortessa X20 flow cytometer equipped with FACSDiva™ software (BD Biosciences). Data analysis was performed using FlowJo™ software (TreeStar, Inc.). At least 0.5×10⁶ events per sample were analyzed to assess CD69 expression and intracellular IFN-γ production.

#### Neutralization assay

Single-round neutralization assays in Tzm-bl target cells (NIH AIDS Reagent Program) (56) were run as previously described (57, 58)}. The antibody VRC07-523LS was provided by the Vaccine Research Center (VRC), National Institute of Health (NIH). Input virus dilution of stocks were calculated from titration experiments to ensure sufficient luciferase output within the linear portion of the titration curve (45,000 RLUs) and were run in the presence of 3.5 μM Indinavir (Sigma-Aldrich) to prevent viral replication. 10 μl of five-fold serially diluted mAbs from a starting concentration of 50 μg/ml were incubated with 40μl of virus in duplicate for 30 minutes at 37°C in 96-well clear flat-bottom black culture plates (Greiner Bio-One). TZM-bl cells were added at a concentration of 15,000 cells per 20μl to each well in DMEM containing 20 μg/ml DEAE-dextran (Sigma-Aldrich). Cell only and virus only controls were included on each plate. Plates were incubated for 24 hours at 37°C in a 5% CO_2_ incubator, after which the volume of culture medium was adjusted to 200 μl by adding complete DMEM. 48 hours post-infection, plates were read by removing 100 μl of media from each well and adding 100 μl of SpectraMax Glo Steady-Luc reporter assay reagent (Molecular Devices) to the cells. After a 15-min incubation at room temperature to allow cell lysis, the luminescence intensity was measured using a SpectraMax i3x multi-mode detection platform per the manufacturers’ instructions. Neutralization curves were calculated by averaging duplicate wells and comparing luciferase units of wells containing antibody to virus-only controls after background subtraction. Curves are fit by nonlinear regression using the asymmetric five-parameter logistic equation in Prism 9 for macOS (GraphPad Software, LLC). The 50% and 80% inhibitory concentrations (IC_50_ and IC_80_) are estimates of the antibody concentrations required to inhibit infection by 50% and 80%, respectively.

#### Co-culture Experiments

A subset of PBMCs were rested overnight, while the remaining cells were frozen for future NK cell isolation. Naïve CD4+ T cells were isolated from the rested PBMCs using the EasySep™ Human Naïve CD4+ T Cell Enrichment Kit (STEMCELL Technologies, cat# 19555). These cells were activated at a concentration of 0.5×10⁶ cells/mL with αCD3/CD28 Dynabeads (1:1 bead to cell ratio) in complete RPMI medium supplemented with1μg/mL of αIL-4,200 μg/mL of αIL-12, and 50 μg/mL of TGF-β for three days. On day 3, beads were removed, and the cells were cultured with 30 IU IL-2 in complete RPMI until day 5. On day 7, the CD4T cells were infected with the HIV-1 NL4-3, JR-CSF, Z331F,190049, MJ4, and 191947 by spinoculation. On day 10, the cells were plated at a high density in 96-well round-bottom plates to promote viral spread and maximize infection efficiency. At day 11, frozen PBMCs were thawed, and NK cells were isolated using the EasySep™ Human NK Cell Isolation Kit (STEMCELL Technologies, cat# 17955). NK cells were rested overnight and treated with or without a TAPI-1 inhibitor in the presence or absence of IL-12, IL-15, and IL-18.

On day 12, CD4T cells were adjusted to the desired density and co-cultured with autologous NK cells at a 1:1 effector-to-target ratio. Antiretroviral therapy (ART) consisting of 1 μM Raltegravir and 500 nM Nelfinavir was included to limit viral replication. For natural cytotoxicity assays, NK cells were cultured as described. ADCC experiments were performed in the presence of 1 μg/mL various bNAbs, such as N6, VRC07-523-LS, VRC07-523-GRLR, VRC07-523-LPLIL, VRC07-523-GASDALIE, or VRC07-523-AAA, with or without IL-12, IL-15, and IL-18 pre-treatment of NK cells at the same concentrations as described above. After a 24-hour incubation at 37°C, cells were prepared for flow cytometric analysis to measure HIV infection. At least 2×10^5^ cells were stained. The staining process involved washing cells with FACS buffer (PBS containing 2% FBS and 2 mM EDTA) followed by staining with eBioscience™ Fixable Viability Dye eFluor™ 450 (Thermo Fisher Scientific, cat# 65-0863-18) at a 1:100 dilution for 10 minutes at 4°C. After washing, human BD Fc Block (BD Biosciences, cat# 564220) was added, and cells were incubated for 10 minutes at room temperature. Surface markers were stained using antibodies at 1:100 dilutions in FACS buffer, including anti-human CD4 APC (Thermo Fisher, cat# 17-0048-42), anti-human CD3 BV786 (BD Biosciences, cat#563918), anti-human CD56 PerCP-Cy5.5 (BioLegend, cat# 362506). Cells were fixed and permeabilized using Cytofix/Cytoperm (BD Biosciences, cat# 554722) and stained intracellularly with p24 FITC (Beckman Coulter, cat# 6604665) diluted in 1× Perm Wash buffer. After a 30-minute incubation at 4°C, cells were washed and resuspended in FACS buffer for analysis.

Percent Remaining p24^+^ cells was determined by comparing the percentage of infected cells in co-culture to the percentage in the control (media only) using the formula:

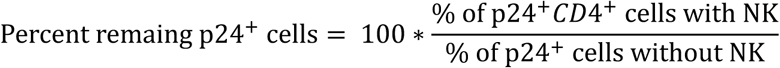

ADCC was calculated by comparing the percentage of NK cells with no antibody, NK cells with antibody and the percentage of infected cells in the control as previously shown (59).

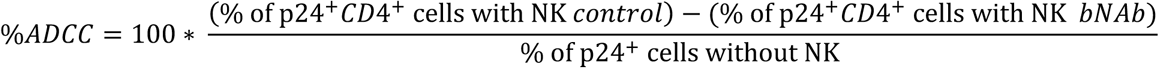

#### Degranulation staining

Co-cultures were performed in the presence of anti-CD107a PE (Southern Biotech, cat#9835-09) at 1:100 dilution and incubated for 4-hour at 37°C. Cells were then washed in PBS and incubated with Human Fc Block (BD Biosciences, cat# 564220) for 10 minutes at room temperature. Following, cultures were stained using antibodies at 1:100 dilutions in FACS buffer, including, anti-human CD3 BV786 (BD Biosciences, cat#563918), anti-human CD56 PerCP-Cy5.5 (BioLegend, cat# 362506) and eBioscience™ Fixable Viability Dye eFluor™ 450 (Thermo Fisher Scientific, cat# 65-0863-18). Staining was conducted for 30 minutes at 4°C and cells were fixed 3% PFA.

#### Envelope staining

Broadly neutralizing antibodies were conjugated with Alexa fluorophore using the Zenon^TM^ Alexa Fluor^TM^ 488 Human IgG Labeling Kit (Invitrogen), following the manufacturer’s protocol. Infected cells were then stained with extracellular anti-CD4 APC (Thermo Fisher, cat# 17-0048-42) and conjugated Env antibodies, followed by the intracellular p24 PE (Beckman Coulter, cat# 6604667).

#### HLA-ABC staining

Infected CD4 T cells were stained with extracellular anti-CD4 FITC (BioLegend, cat# 300506) and anti-HLA-ABC (BioLegend, cat# 311409), followed by the intracellular p24 PE (Beckman Coulter, cat# 6604667).

#### Clustering analysis

Hierarchical clustering was performed using ClustVis (BETA) (https://biit.cs.ut.ee/clustvis/). The analysis was based on the mean percentage of remaining p24⁺ cells or ADCC. Data were used without transformation, with row centering and row scaling both disabled. Rows were clustered using the Euclidean distance metric and the average linkage method, with tree ordering based on the tightest cluster first. Columns were also clustered using Euclidean distance and the average linkage method, with tree ordering set to display the tightest cluster first.

#### Statistics

Flow cytometry data were analyzed using FlowJo software (BD), and all statistical analyses were performed in GraphPad Prism version 10.2.1 (464). Differences between treatment conditions and donor groups were assessed using two-way ANOVA with multiple comparisons, unless otherwise specified. Friedman test with multiple comparisons was used to evaluate the activation of the VRC07-523-LS and its mutant compared to no bNAbs. Friedman test with multiple comparisons, was used to evaluate significance of cytokine secretion between no bNAbs to WT and mutants. Wilcoxon signed-rank test was used to evaluate differences within conditions for the percentage of p24⁺ cells (tested against a hypothetical value of 100) and for ADCC (tested against a hypothetical value of 0), as well as to assess interactions using the Bliss model of synergy (tested against a hypothetical value of 0). Envelope staining and the percentage of remaining p24⁺ cells were analyzed using two-way ANOVA. Comparisons between DMSO and TAPI-1 treated conditions were performed using paired t tests. The Wilcoxon signed-rank test was also applied to compare no cytokine versus cytokine conditions under both untreated and TAPI-1 pretreated settings. One-way ANOVA was used for analyses of IFN-γ production, and differences between male and female donors were determined by two-way ANOVA.

Spearman correlation analyses were used to evaluate associations between CD16 expression and CD69 induction, Env binding and neutralization IC_50_ values, ADCC activity and surface Env binding, IFN-γ fold change and CD16 gMFI fold change, as well as between CD69 gMFI and donor age. Pearson correlation was used to assess the relationship between the percentage of remaining p24⁺ cells and HLA-ABC fold change. P values < 0.05 were considered statistically significant.

#### Study Approval

White blood cell concentrates (buffy coat), derived from a single unit of whole blood through centrifugation, were obtained from the Gulf Coast Regional Blood Center. Blood donors were volunteers aged 17 and older. Only age and biological sex information were collected; no other personal details were provided.

#### Data Availability

The data points presented in the graphs are available in the Supporting Data Values file. Additional data can be found in the supplement.

### Author contributions

C.M. designed and performed experiments, analyzed data, and wrote the manuscript. A.B. supervised the study, contributed to conceptualization and design, and acquired funding. R.M.L. and T.M. generated and provided the bNAb mutants and contributed to discussions on antibody selection and experimental design. M.K. performed and analyzed the neutralization assays with VRC07-523LS across viral subtypes. J.L. generated and analyzed HLA-ABC data. C.S.H. and E.K.M. provided technical support and assisted with experimental execution. All authors discussed the results, contributed to manuscript revision, and approved the final version

### Funding support

C.M. was supported by NIH NIAID-funded Grant Number T32AI158105 and F31AI186614. Research reported on this publication was supported by the NIAID of the NIH under the grant R21-AI172042 and UM1-AI164565 to A.B. and R.L. This research has been facilitated by the services and resources provided by District of Columbia, Center for AIDS research, an NIH funded program (AI117970), which is supported by the following NIH co-funding and participating institutes and centers: NIAID, NCI, NICJD, NHLBI, NIDA, NINH, NIA, FIC, NIGGIS and NDDK and OAR. The content is solely the responsibility of the authors and does not necessarily represent the official views of the NIH.

## Acknowledgments

C.M. thanks all previous lab members Callie Levinger, J. Nathalie Howard and Amanda Macedo.

